# On Pectin Methyl-esterification: Implications for *In vitro* and *In vivo* Viscoelasticity

**DOI:** 10.1101/565614

**Authors:** J.L. Kaplan, T.A. Torode, F. Bou Daher, S.A. Braybrook

## Abstract

Pectin is a major component of the primary plant cell wall and is important for cell expansion. However, the relationship between its chemistry and mechanical properties is not fully understood, especially *in vivo*. In this study, a protocol for viscoelastic micro-indentation using atomic force microscopy (AFM) was developed and applied to pectin *in vitro* and *in vivo*. After determining that linear viscoelasticity was a suitable theoretical framework for *in vitro* pectin analyses were conducted with both a standard linear solid and fractional Zener model. These indicated a strong coupling between elastic and viscous properties over a range of degrees of methyl-esterification (DM). Both elasticity and viscosity were found to vary non-linearly with DM which had interesting consequences for pectin gels of mixed DM. In *Arabidopsis* cell walls, the standard linear solid model was found to be appropriate. In this *in vivo* composite material a weaker elastic-viscous coupling was exhibited, correlated with DM. The viscoelastic testing *in vivo* of rapidly elongating cell walls, rich in high DM pectin, displayed a longer viscous time-scale. The implications of the testing method and results are discussed in the context of mechanobiology, mechano-chemistry, and cell growth.

## 1 Introduction

Networks of hydrophilic polymer chains, known as hydrogels, are a critically important class of material in biology [1]. Hydrogels made from pectin, and insights into their mechanical properties, are useful in applied [2, 3] and fundamental contexts [4, 5]. In the plant species *Arabidopsis thaliana*, pectin is the single largest constituent of the primary cell wall [6], the structural layer encasing growing cells which is acknowledged to be a critical arbiter of growth [7]. The umbrella term ‘pectin’ refers to a variety of pectic polysaccharides which cohabit the cell wall including homogalacturonan (HG), rhamnogalacturonan I, rhamnogalacturonan II and xylogalacturonan [8]. In *Arabidopsis*, HG is the largest component by a significant margin [9]. Key features of HG are its initial state, which is highly methyl-esterified, and its de-esterification when acted upon by pectin methylesterase (PME) enzymes [10]. HG de-esterification can occur in a blockwise contiguous mode or a non-contiguous random mode. The degree of ‘blockiness’ likely depends on the properties of the PMEs themselves [11], of which there are sixty-six (66) in *Arabidopsis* [12]. Random de-esterification can occur through non-enzymatic means with changes in pH [13]. When a methyl group is removed (i.e. de-esterified) a negative charge is left in its place and two negatively charged sugars can bond using Ca^2+^ ions [11]. Indeed, calcium cross-linking is the primary method of gelation for low degree of methyl-esterification (DM) gels and longer ‘blocks’ of de-esterified units form stronger bonds – provided there is sufficient free calcium nearby. In contrast, high DM HG relies on hydrogen bonds and hydrophobic interactions, localised at the methylated groups, for gelation [3, 14]. In addition to the above summary of pure HG gel mechano-chemistry, it is important to be aware of added complexities *in vivo*. The mechanical integrity of low DM HG can be degraded by polygalacturonases leading to altered gelation and thus cell growth [11,15]. Additionally, more random de-esterification has been linked to cell separation indicating a role beyond gelation alone [16]. Importantly, the pectic polysaccharides in the cell wall coexist with hemicelluloses and cellulose thereby forming a composite material. Evidence suggests that the mechanical properties of *in vivo* pectin are carefully spatio-temporally regulated by plants and thereby play a crucial role in morphogenesis – the process of developing form [4]. Pectin methyl-esterification state has been linked to cell wall elasticity and cell expansion in elongating hypocotyls [17] (an organ whose rapid axial elongation is essential for emergence of the young seedling from the soil). However, there remain a number of open questions regarding the relationship between pectin biochemistry and rheology, and the exact role that pectin rheology plays during cell growth and other developmental processes [17–19].

It has been known for some time that pectin exhibits viscoelastic behaviour *in vitro* [20]. Previous viscoelastic studies of pectin have used dynamic mechanical analysis, in which an oscillatory force is applied, to extract a complex relaxation modulus *G\** [20–25]. Though informative, the absence of a modelling approach in such studies places limitations on what can be inferred from the data and our ability to link rheological behaviour to physical mechanisms of deformation. Only a smaller number of early pectin studies used a model fitting approach [26–29]. In order to bridge the complexity between pure pectin gels and primary cell walls, some have investigated artificial composites containing only selected cell wall components. For relatively large compression of cellulose/pectin mixtures, the presence of pectin was found to increase resistance to compression, possibly due to reduced permeability [30]; viscoelastic properties and DM of the pectin used were found to play a role [31]. A recent study investigating the viscoelastic properties of cellulose/pectin composites found that both the storage and loss modulus decreased approximately 5-fold from DM69 to DM33 [2] – possibly a consequence of insufficient free calcium. Moving to plants, elastic tests and measures of ‘extensibility’ (which has several different meanings [32]) dominate the literature. In particular, AFM-based nano-indentation has been an important tool for localised *in vivo* mechanical testing [33]. However, more recently there has been growing awareness of the cell wall’s viscoelastic nature, which coincides with one of the definitions of extensibility [32]. Viscoelastic investigations have been conducted on *Arabidopsis* organ initiation [18], leaf samples [34], living roots [35], and tomato mesocarp cells [36].

In this study we developed an AFM-based viscoelastic methodology capable of measuring time-dependent mechanical properties of pectin gels and *Arabidopsis* hypocotyl cells at micron indentation depths. This method was used to gather force relaxation data from HG gels and identify linear viscoelasticity as a suitable theoretical framework. A fractional Zener (FZ) model was then identified as an ideal constitutive model due to its ability to capture power-law type behaviour. This model, in conjunction with a more intuitive 2 time-scale standard linear solid (SLS2), were used to identify ionic bond reformation as the dominant mode of physical deformation in gels under these types of load. The models were further used to investigate mixed DM HG gels. Our recent work [17] showed that in the elongating dark-grown *Arabidopsis* hypocotyl, epidermal cells display an elastic asymmetry coincident with a difference in pectin DM; faster growing axial cell walls were more compliant and had a higher DM compared to slower growing transverse walls. As the hypocotyl system provided us with an ideal testing opportunity for our AFM-based micro-creep method we decided to build on the previous elastic study. Short time-scale results mirrored previous elastic work closely but at longer time-scales the compliance of axial and transverse walls appeared to get closer, suggesting a possible coupling may be at work.

## 2 Materials and Methods

### 2.1 Preparation of HG Gels

Homogalacturonan samples (Herbstreith & Fox, UK) of varying degrees of methyl-esterification (DM) (33, 41, 50, 60 and 70; or blends thereof) were dissolved in de-ionised water, neutralised with NaOH to pH6.5, and made up to 1.5% (w/v). The HG Samples had the following galacturonic acid contents, as supplied by Herbstreith & Fox: DM33 - 84%, DM41 - 90%,DM50 - 86%, DM60 - 89%, DM70 - 83%); the samples were therefore mostly HG but not completely pure. Homogalacturonan gels were set using the CaCO_3_/gluconolactone (GDL) method commonly used to set calcium cross-linked hydrogels [37], via addition of CaCO_3_ (Cat. No. 21061, Sigma, UK) and GDL (Cat. No. G4750, Sigma, UK) into the pectin solution to facilitate the controlled, slow release of Ca^2+^ ions. The pectic/CaCO_3_/GDL mixture was left on a microscope slide for 12 hours at room temperature to fully set prior to use in experiments. The ratio of calcium ions available to bind pairs of de-methylated galacturonan residues (*R*), was kept at *R* = 1 for all gels. *R* = 2 × [Ca^2+^]/[COO ^−^]. A stoichiometric ratio of GDL, [GDL] = 2 × [Ca^2+^], was used to solubilise all CaCO_3_ in the gel, without altering the pH via excessive proton release. Prior to AFM experiments, the homogalacturonan hydrogels were soaked for 30 minutes in CaCl_2_ solutions of equal molarity of Ca^2+^ as within the hydrogels.

### 2.2 Preparation of Plant Material

*Arabidopsis thaliana* ecotype Col-0 seeds were surface sterilized with 70% ethanol for five minutes then rinsed with 100% ethanol for one minute and left to dry prior to plating them on half strength Murashige and Skoog (MS) media. Seeds were stratified for 2 days at 4 ^*◦*^C in the dark before transferring them to a growth chamber with 16/8-hours light/dark cycle at 20 ^*◦*^C for germination. Germinated seeds were transferred to half strength MS plates containing 87.6 mM sorbitol, sealed with aluminum foil and left to grow for 24 hours prior to AFM scanning.

### 2.3 AFM-based indentation tests

#### 2.3.1 Gels: Elastic and Viscoelastic Tests

A 50 µm diameter silicon bead (Cat# C-SIO-50.0 from Microspheres-Nanospheres, Corpuscular, Cold Spring, NY, USA) was mounted on a 2.8 N m^−1^ tipless cantilever (TL-FM-10, Windsor Scientific Co., UK) for DM50 gels and DM50 mixtures, and a 48 N m^−1^ tipless cantilever (TL-NCL-10, Windsor Scientific Co., UK) for lower DM gels, using standard 2-component epoxy as in [33]. After the gels had been left to swell for 30 minutes in CaCl_2_ (of equal molarity of calcium ions as within the hydrogels) the slides were carefully placed on the stage of a Nano Wizard 3 AFM (JPK Instruments, DE) and covered with just enough CaCl_2_ solution to cover the gels. For elastic tests, a new (previously unindented) point near the centre of the gel was then subjected to a range of elastic indentations over a force range that was found to contain the indentation depths of interest. At each point the up/down direction of force variation was alternated to ensure pre-stress was not affecting the results (no pre-stress effects were observed). For viscoelastic indentations, the same settings as for elastic were used apart from the introduction of a 15 second constant height hold. All indentations, both elastic and viscoelastic, were done at a fast indentation rate of 50 µm s^−1^.

#### 2.3.2 Plants: Elastic and Viscoelastic Tests

24 hour old seedlings were gently stuck on a double sided tape placed on a glass slide, immediately covered with a solution of 600 mM mannitol and plasmolysed for 15 minutes prior to scanning. A Nano Wizard 3 AFM (JPK Instruments, DE) mounted with a 10 nm diameter pyramidal tip (Windsor Scientific, UK) on a 45 N m^−1^ stiffness cantilever was used for indentations. For elasticity testing, an area of 100 µm × 50 µm at the base of the hypocotyl was indented with a resolution of 64 × 32 points using 500 nN force. Indentation speed used was 100 µm s^−1^. For viscosity testing, 10 points were chosen on the cell wall of a basal cell (5 on the transverse wall and 5 on the axial wall) with a 15 second indentation hold time on each point. Seven hypocotyls were used. For viscoelastic tests 19 transverse and 18 axial wall indentation points were analyzed from across these seven hypocotyls. For elastic analysis 19 transverse and 15 axial wall indentation points were used.

### 2.4 Mechanical Testing

Within the range of depths tested, linear elastic and linear viscoelastic theory was found to be appropriate by the methods discussed in the supplementary information.

#### 2.4.1 Hertz Contact Model for A Spherical Indenter

The Hertz contact model for a rigid sphere indenting a flat elastic half-space modified appropriately for AFM indentation [38] is

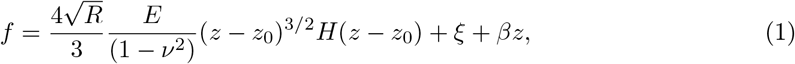

where *f* is force applied, *R* is the radius of the indenting sphere, *E* is the Young’s Modulus, *ν* is the Poisson ratio which is assumed to be 0.5 (implying incompressibility), *z* is the position of the indenter as represented internally by the AFM, *z*_0_ is the reference contact point, *H* is the Heaviside step function, *ξ* and *β* are used to represent the offset and drift respectively that commonly occur during AFM indentation. For elastic analysis, the offset and drift were found to be minuscule for the majority of files but in any case were subtracted from the data using the JPKSPM data processing software provided by the AFM manufacturer. For viscoelastic analysis *β* was taken as 0 in order to not affect the constant deformation/force section of the data for relaxation/creep tests respectively. The reference contact point *z*_0_ is not known *a priori*. Various methods of identifying the contact point have been proposed [38–40]. In this study, the contact point was found using a qualitatively determined force threshold. A specific threshold was selected for each material tested by plotting the force/displacement curves for that material semi-logarithmically in the force axis and observing where the largest change in force took place – which corresponds to initial contact. After this the material displacement, *δ*, could be found using the relation *δ* = *z −z*_0_.

For viscoelastic testing the elastic-viscoelastic correspondence principle [41] can be used to derive the time-dependent analogue of Equation 1 for displacement controlled (relaxation) tests. Noting that *E* = 3*G* for the incompressible case, where *G* is the shear modulus, the linear viscoelastic relaxation equation is

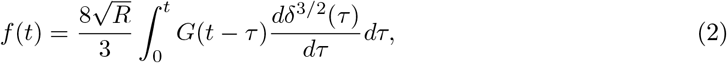

where *f*, *δ* and *G* now represent dynamic functions and *τ* serves as an integration dummy variable. A similar method can be followed [42] to derive the dynamic force-displacement relationship for stress controlled creep tests which results in

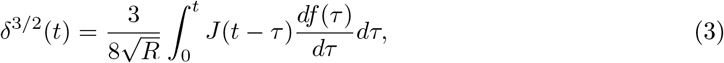

where *J* is the material creep function and all other symbols have their previous meaning. For ramp and hold loading, the above integral can be simplified using a ramp correction factor [41, 42]. However, the complex force/displacement feedback cycle used by the AFM meant it was not feasible to generate an accurate linear ramp. Instead, a convolution was performed using the NumPy Python package (numpy.convolve), thus enabling arbitrary loading kernels to be used.

#### 2.4.2 Standard and Fractional Viscoelastic Models

The relaxation and creep moduli of the SLS2 model take the form of Prony series:

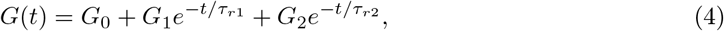

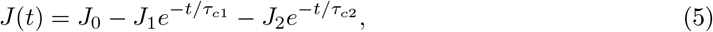

where *τ_r_* and *τ_c_* are relaxation and creep (retardation) time-scales respectively. Elastic/viscous ratios can be defined on the relaxation [41] and creep moduli in the following way:

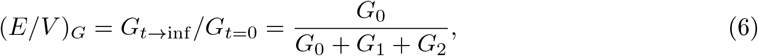

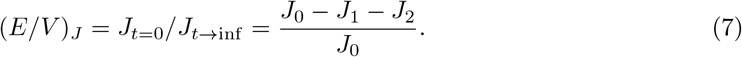

The FZ model is a combination of an elastic spring in series with a fractional Kelvin-Voigt element (1 spring and 1 spring-pot in parallel). The relaxation shear modulus and creep shear compliance of the FZ model can be written [43]:

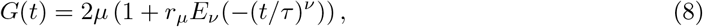

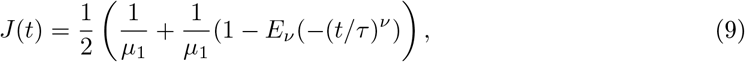

where *E_ν_* is shorthand for *E_ν,_*_1_ – the Mittag-Leffler function with the second parameter equal to 1 and the first parameter, the spring-pot parameter, allowed to vary between 0 and 1 reflecting the fact that a spring-pot can behave as a spring (*ν* = 0) or a dash-pot (*ν* = 1). All other parameters are derived model parameters.

### 2.5 Statistical analysis

When statistical tests were necessary a Wilcoxon Rank-Sum Test was performed in Python (scipy.stats.ranksums). This non-parametric test performs well with samples of unequal variance and non-normal distributions.

## 3 Results and Discussion

### 3.1 Fractional Zener Viscoelastic Model Provides the Best Representation of Pectin Gel Behavior

The fitting of viscoelastic models to data is helpful for quantitative comparisons of viscoelastic materials. Standard viscoelastic models are often represented schematically using a combination of springs and dash-pots, resulting in creep or relaxation moduli that take the form of a sum of exponentials. More recently, there has been increasing interest in the use of spring-pots – a viscoelastic element derived using fractional calculus. For example, the recent paper by Bonfanti et al. which identified a fractional viscoelastic model that was able to capture and predict the viscoelastic behaviour of epithelial monolayers, as well as individual cells [44]. This generalised element allows for the spectrum of behaviours in-between a spring and a dash-pot to be explored. In isolation, the spring-pot exhibits power-law behaviour. In combination with springs and dash-pots it is able to accurately capture complex combinations of exponential and power-law viscoelasticity.

To make an informed decision on which model to use, the qualitative behaviour of the gels’ relaxation was examined. It can be seen from the averaged DM41 data (*N* = 20 across 6 gel samples) in Figure 1a, that the gels exhibited a brief period of rapid relaxation followed by power-law relaxation until the end of the hold. Due to the observed complex decay signature, it was hypothesised that a fractional model would be best suited to model the behaviour of HG gels under load. The FZ model was chosen as it is a fractional analogue of the widely used standard linear solid model. One drawback of using fractional viscoelastic models is that physical interpretation of their parameters can be difficult. As such, it was decided that an SLS2 model would also be used, to provide a more holistic insight into the gels behaviour. The averaged data (DM41, 2 µm depth), along with the averaged fits of the two models are shown in Figure 1b. The superior qualitative fit of the FZ model should be noted. Quantitatively, the sum of squared residual error for the SLS2 model was 1.2e–10 and the error for the FZ model was 3.6e–11, an order of magnitude smaller. From a phenomenological perspective, the suitability of a fractional model may be justified by considering the heterogeneous distribution of calcium cross-linked block size and density within pectin gels that may yield a large number of inherent material time-scales – a feature that is known to give rise to power-law type viscoelasticity [45]. The modelling approach to quantifying pectin rheology is in contrast to some previous works which have taken a model-free approach via comparison of the experimental complex modulus *G\** over a range of tested frequencies [2, 20, 22, 23, 25]. However, this model-free approach places limits on the amount of physical insight that can be gained from the data. Although some of the earlier pectin studies did use a modelling approach based on springs and dashpots [26, 27], they often featured a large number of parameters which can lead to overfitting [28, 29]. The use of these complicated models may have been motivated by the appearance of a better fit, given that a large number of spring and dashpots can approximate power law behaviour [46]. However, as noted by others [44, 46], the fractional springpot element allows the fitting of power law behaviour with much lower risk of excessively complex model selection, thus facilitating scientific parsimony.

**Figure 1:**
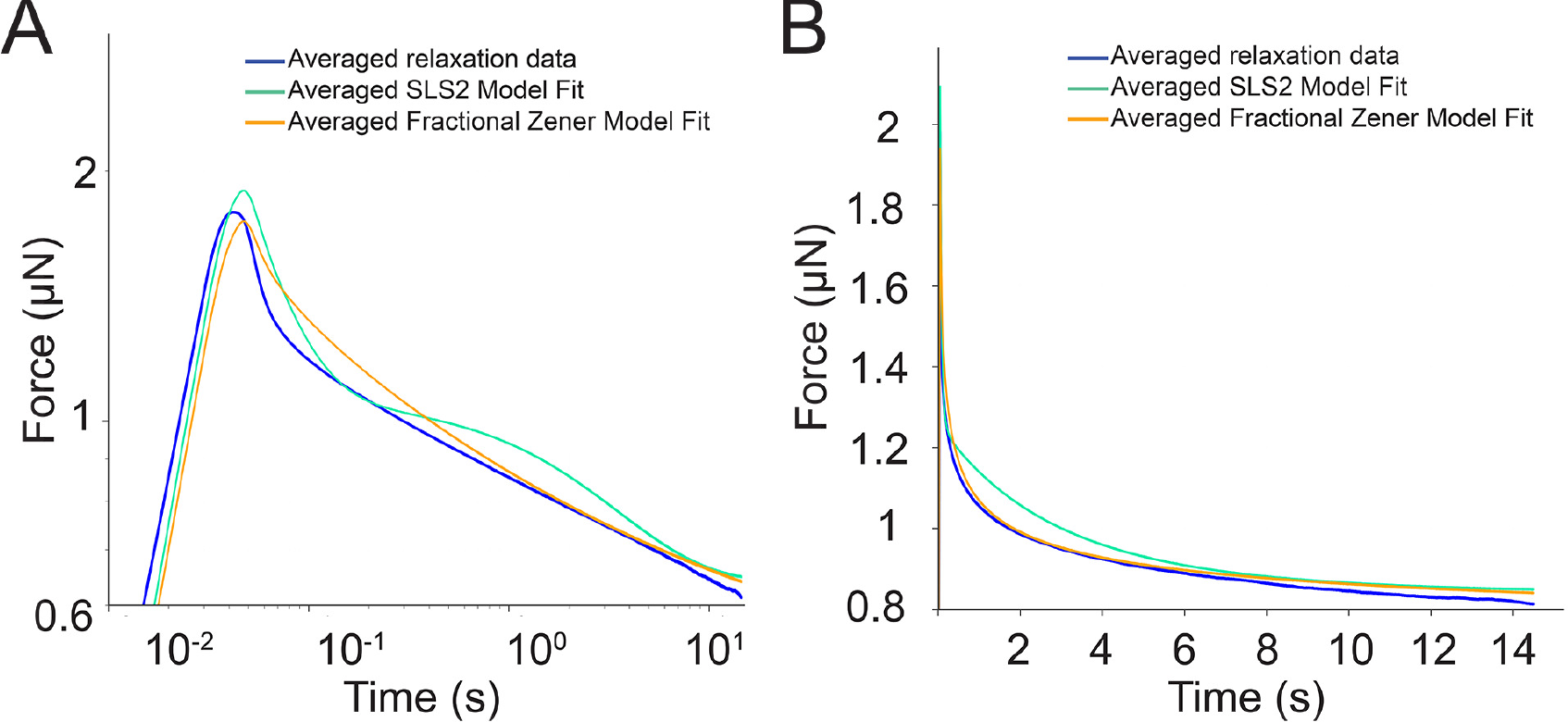
a) Averaged relaxation data from DM41 viscoelastic relaxation tests plotted in log-log scale. b) Averaged relaxation data from DM41 viscoelastic relaxation plotted in normal scale with the predictions from the two tested models also plotted.

### 3.2 Elasticity and Viscosity Are Inversely Correlated with DM

Pectin can be generated with different DMs, which alters the cross-linking behavior and gel properties [4, 37, 47]. Within the plant cell wall, pectin can exhibit differing degrees of DM which contributes to differential wall elasticity [18]. To assess the effect of DM on viscoelastic properties in a minimal system, we performed AFM-based force relaxation tests on HG gels with different DM but the same amount of available calcium (DM 33, 41, 50; *R_eff_* = 1, see Methods). Indentations were performed at forces of 2506 nN, 1911 nN and 276 nN for DM 33, 41 and 50 gels respectively; this allowed for an approximately fixed depth (2 µm) of indentation across all samples. Resulting curves were analysed using both an SLS2 model and an FZ model. The fitting results are shown in Figure 2.

**Figure 2:**
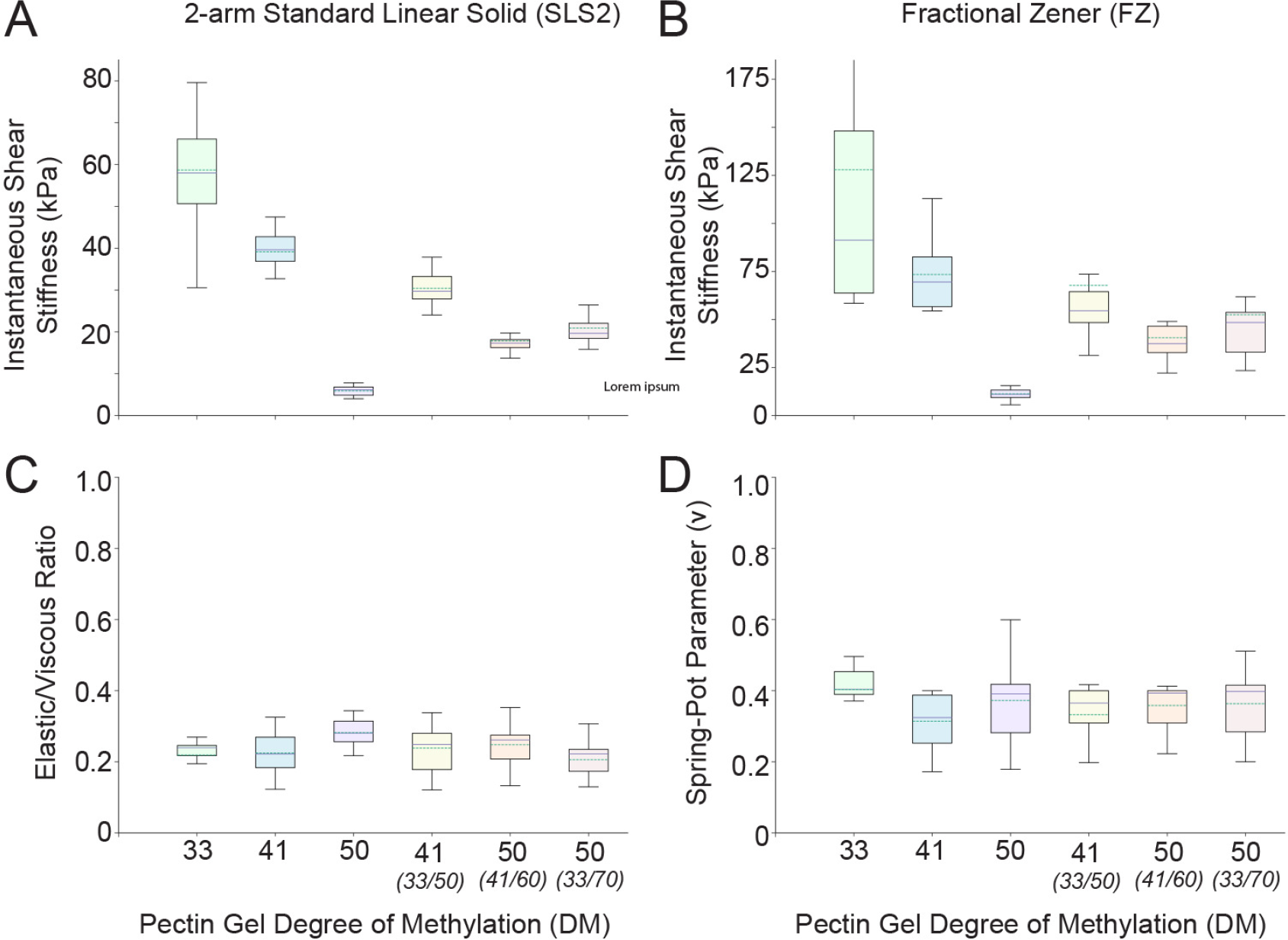
HG model fit results – continuous line represents median of data and dashed line represents mean of data. a) ISS, or equivalent shear stiffness at time t=0, as found by the SLS2 model for the range of tested DM gels. b) ISS, or equivalent shear stiffness at time t=0, as found by the FZ model for the range of tested DM gels. c) Elastic/Viscous ratio as found by the SLS2 model for the range of tested DM gels. d) Spring-pot parameter *ν* as found by the FZ model for the range of tested DM gels.

There was a large difference in the absolute values of instantaneous shear stiffness (ISS) between the two models. Absolute values obtained here may be highly sensitive to model parameters, testing methods and experimental conditions; relative differences were considered to be more informative in this context. Compared to the SLS2 model, the FZ ISS was 118%, 88% and 92% higher on average for DM 33, 41 and 50 respectively. This suggests that the Fractional model attributed more of its resistance to its instantaneous elastic elements whilst the Standard model attributed more of its resistance to its viscous elements (dash-pots). To gain understanding of the gels inherent mechanical properties, we focused on the consistencies between the two models’ results. The relative differences in ISS between gel types follow a similar qualitative trend for both models. Both showed a slight decrease in ISS in DM 41 pectin compared to DM 33, and a significant decrease in ISS at DM 50. The Elastic/Viscous ratios shown in Figure 2c imply that viscous effects varied proportionally to elastic effects in the SLS2 model. Similarly, Figure 2d shows that the *ν* parameter of the FZ model – which determines whether the spring-pot behaves more like a spring or a dash-pot – remained fairly consistent between gels tested. As there is only one spring-pot and no dash-pots in the FZ model, this parameter *ν* is the sole determinant of relative elastic or viscous behavioural dominance.

The results discussed above indicate two important points regarding pectin gel mechanical properties and their relationship to DM. Firstly, irrespective of the viscoelastic model, there was a clear inverse relationship between DM and gel strength (both elasticity and viscosity). This result is consistent with previous rheological tests [16, 22, 48, 49], and conforms to the idea that – assuming a sufficient degree of blockiness – the lower the DM the higher the number of available junctions for forming inter-chain bonds with available calcium ions. Secondly, irrespective of the viscoelastic model, there is a tight coupling between elastic and viscous parameters across different DMs. This coupling has been found in previous studies as well [22, 23]. In particular, Fraeye et al. [22] performed small deformation oscillatory tests on a large variety of pectins over a range of DM, calcium concentration (R value), degree of blockiness, and frequency and the coupling between the storage modulus and experimental modulus was consistently striking. This coupling suggested a single unified physical deformation and dissipation mechanism that varies with DM and determines both short and long time behaviour. The simplest candidate for this mechanism is the breaking and reforming of ionic chains as it would successfully explain both the inverse gel strength relationship with DM, and the tight coupling of elastic and viscous properties. Interestingly, this was the same conclusion reached by Zhao et al. [50] in their rheological study of pectin’s algal equivalent, alginate, when the alginate was gelled ionically. Finally, it should be noted that this dominant mode of deformation supports the hypothesis put forward in the previous section – that a power-law relaxation is observed due a heterogeneous distribution of ionic chain lengths breaking and reforming.

### 3.3 Pectin Strength vs. DM is Non-Linear – Consequences For Mixed DM Gels

A clear non-linear relationship between pectin DM and gel strength was observed. (As elastic and viscous properties were found to be tightly coupled, the qualitative term ‘strength’ will henceforth be used to refer to their aggregate value.) The difference in gel strength between DM 41 and DM 50 was far greater than the difference between DM 33 and DM 41 (50% and 564% difference respectively, when comparing mean ISS from the SLS2). This may be due to the fact that a minimum number of consecutive calcium bonds (approximately 7 - 20) are required to form a stable inter-chain bond [13, 51, 52]. Thus, higher DM gels may struggle to attain the minimum number of consecutive bonds required for stable bond formation. Slightly below DM 50, there may exist a sharp DM threshold above which strong, stable gels do not form. Initial tests were attempted on higher DM (60 and 70) gels but they were found to be so soft as to be incomparable to lower DM gels at similar indentation depths. To use the language of the Flory-Stockmayer percolation model [53], this DM threshold may be analogous to a ‘percolation threshold’ which is a type of gel/solution transition point.

In addition to the above, we were also interested in testing the properties of mixed DM pectin gels. The reason for this is that plant cell walls likely contain heterogeneous DM pectin. We generated two mixed gels with an effective DM 50, one using a combination of DM 33 and DM 70, and another using a combination of DM 41 and DM 60. We also generated one mixed gel with an effective DM 41 using a combination of DM 33 and DM 50 pectin. These were subjected to the same AFM indentation tests and model fits as above to investigate whether their mechanical properties would be similar to the homogeneous DM pectin gels. As in the above case, the elastic and viscous properties were found to be coupled across gel types, as shown in Figures 2c and 2d. In contrast to the above, the strength of the gel mixtures was found to be significantly different to their effective homogeneous DM counterpart. Comparing SLS2 ISS (though the trend was the same in both models): the effective DM 41 was 22% weaker than pure DM 41, the effective DM 50 (33/70 mix) was 255% stronger, and the effective DM 50 (41/60 mix) was 202% stronger than the pure DM 50 gel. Similar results were found by Löfgren and Hermansson [24], who also showed that the fine structure of high/low DM mixtures were heterogeneous in architecture, with clusters of low DM pectin forming (though this heterogeneity is unlikely to have affected our results as the length-scale of heterogeneity appears far smaller than our experimental length scale of 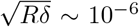). These differences may be explained by the non-linear relationship between pectin DM and mechanical strength discussed above. Given that pure DM 50 ISS is an order of magnitude less strong than DM 41 and DM 33, and DM 60 and DM 70 have almost negligible strength in comparison to those two gels, it makes sense that a DM volume fraction mix of the 41/60 or 33/70 would be significantly stronger than DM 50 alone. (Indeed, the fact that the DM 50 mixtures’ mechanical strength is dominated by the low DM contribution explains why the DM 50 mixture based on DM 33 was slightly stronger than the DM 50 mixture based on DM 41, as DM 33 is slightly stronger than DM 41.) The reverse phenomenon explains the relative weakness of the effective DM 41 mixture.

### 3.4 The 2 Time-Scale Standard Linear Solid Model is Informative for Plant Cell Wall Creep Data

We have recently shown that the rapidly elongating axial cell walls of the dark-grown *Arabidopsis* hypocotyl epidermis displayed a reduced elastic modulus and increased amount of high DM pectin, when compared to more slowly growing walls in the same sample [17]. As the tests in that study were solely elastic, not viscoelastic, this system provided us with an ideal testing opportunity for our AFM-based rheology method *in vivo*. It also allowed us to investigate whether our rheological results in gels translated to the *Arabidopsis* cell wall, which is comprised of pectin and other polysaccharides such as cellulose. In order to apply a similar methodology as above to *Arabidopsis* cell walls, some adaptations were required. The methodology used above with gels for the evaluation of poroelastic vs. viscoelastic dominance is less appropriate in the context of the primary cell wall; there are a multitude of potential sources of non-linear mechanical behaviour in the plant cell such as turgor pressure, bending stiffness of the wall, and geometric effects due to its cellular-solid structure. As pectin is the largest constituent of the primary cell wall in *Arabidopsis*, it was assumed that the dominant behavior would be viscoelastic as in gels. Although artificial *in vitro* pectin/cellulose composites have been shown to exhibit poroelasticity [30, 31] the experimental length scales in those studies were significantly larger meaning porous effects would have been more dominant, and furthermore it is difficult to extrapolate from pectin/cellulose composites to the vastly more complex cell wall. Creep tests (fixed force, height evolves in time) were performed on the plants, as opposed to force relaxation on gels. There were two main reasons for this: firstly, the AFM signal was found to be far noisier and creep tests provided the best signal/noise ratio due to the force-feedback AFM controller. Secondly, the data would be more comparable to organ level extensometer data in the literature [54].

As with the gels, it was necessary to first identify a suitable model with which to analyse the viscoelastic data. Ideally, the model would also be commensurate with the models used for the gels. Exponential relaxation type models have been successfully used in various mechanical studies of plant cells [34,36]. Due to its ability to capture complex mechanical behaviors the FZ model has been used in several animal cell biomechanical studies [cite epithelial] [55–57] but it is uncommon and has not thus far been used for plant cell rheology. Given that these two models performed best with pectin gels, we chose to test both the SLS2 and FZ models for their suitability in analysing *in vivo* wall viscoelasticity.

First, an elastic stiffness map was generated using the same AFM-based nanoindentation method utilized in the previous study [17,58]. The basal, rapidly elongating, region of the hypocotyl was exposed to rapid indentation within the elastic range of plant tissues, fast enough to negate any viscous behavior [58], and a Hertzian indentation model was used to generate a map of ‘indentation modulus’ (IM) 24 hours post-germination; Figure 3a); indentation modulus is defined here as the Hertz-derived Young’s Modulus (E) for a plant cell where the cellular-solid nature of the material is non-standard for analysis of E. The IM map showed that the transverse walls were significantly stiffer than the axial walls (Transverse: 13.60 ± 1.82 MPa, Axial: 6.19 ± 0.60 MPa, *p* < 0.001), in strong qualitative agreement with our previous work [17]. The results also agree qualitatively with those found in another study [19], although technical differences in experiment and analyses make the absolute values incomparable. The IM map was used to pick points on each sample’s axial and transverse walls (5 points on each wall type per sample) at which creep tests were conducted of duration 15 s and force 500 nN. Creep data was then fitted to both proposed viscoelastic models. It was found that the spring-pot parameter of the FZ model, *ν*, was equal to 0.9958 ± 0.0031 and 0.999 91 ± 0.000 01 for axial and transverse walls respectively. This implied that the fractional spring-pot element was behaving as a standard dash-pot and that any power-law growth in compliance was negligible. This fits with the experience others have had using 1 and 2 time-scale SLS models in plant systems [34, 36]. Further, in our early work we had effectively used a Kelvin-Voigt model in preliminary viscoelastic analysis of plant cell tissues [18] which although simpler than the SLS, is still based on a decaying exponential with a well defined time scale. Based on the above data and context, we concluded that use of the SLS2 model was sufficient for further analysis; a representative fit is shown in Figure 3b.

**Figure 3:**
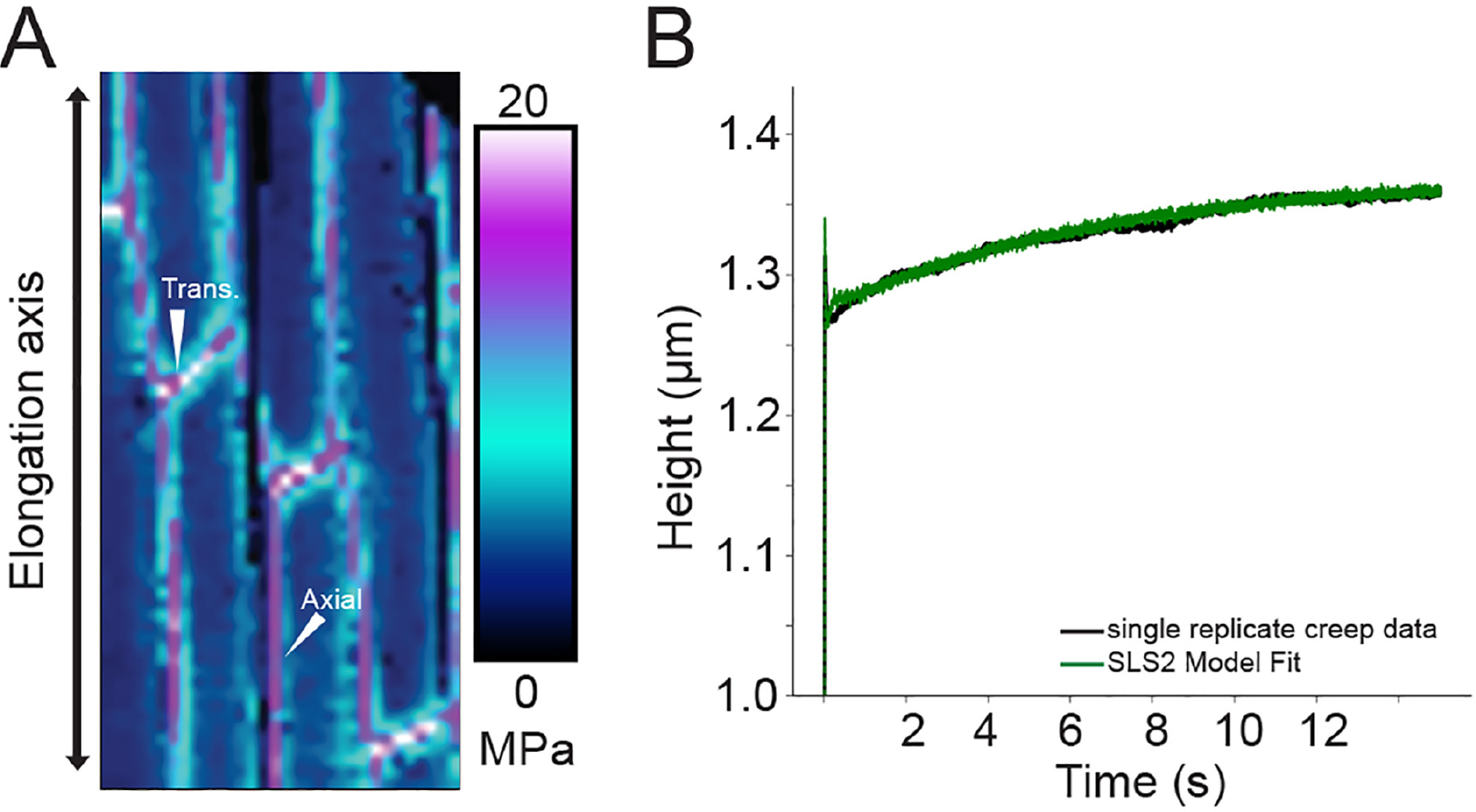
a) Representative map of elastic indentation moduli over a section of *Arabidopsis* hypocotyl. b) Representative *Arabidopsis* hypocotyl viscoelastic creep data with SLS2 fit.

### 3.5 *In vivo* DM Negatively Correlated With ISS and Rate of Creep – Predicted Plateau Shear Stiffness Appears To Be Coupled In Axial/Transverse Walls

Since we knew that transverse walls exhibited lower DM [17], we could explore how DM was related to the SLS2 parameters in a more complex wall material. Figure 4a shows the ISS of the SLS2 model for both transverse and axial cell walls. As with the elastic force map, the ISS data demonstrate that at near-instantaneous time-scales the axial walls are significantly less stiff than the transverse walls (315 ± 46 kPa for axial walls compared to 488 ± 104 kPa for transverse walls, a 55% difference, *p* < 0.05). This difference is in line with the elastic results found in this (Figure 3a), and earlier studies [17, 19]. When combined with our gel data above our results fit the hypothesis expounded by Bou Daher et al. [17] that lower DM pectin in the transverse hypocotyl cell wall results in an increased stiffness compared to the axial wall. However, it is interesting that the ISS values found by fitting the viscoelastic SLS2 model are approximately one order of magnitude smaller than the IM values at the same point, found using an elastic Hertz fit.

**Figure 4:**
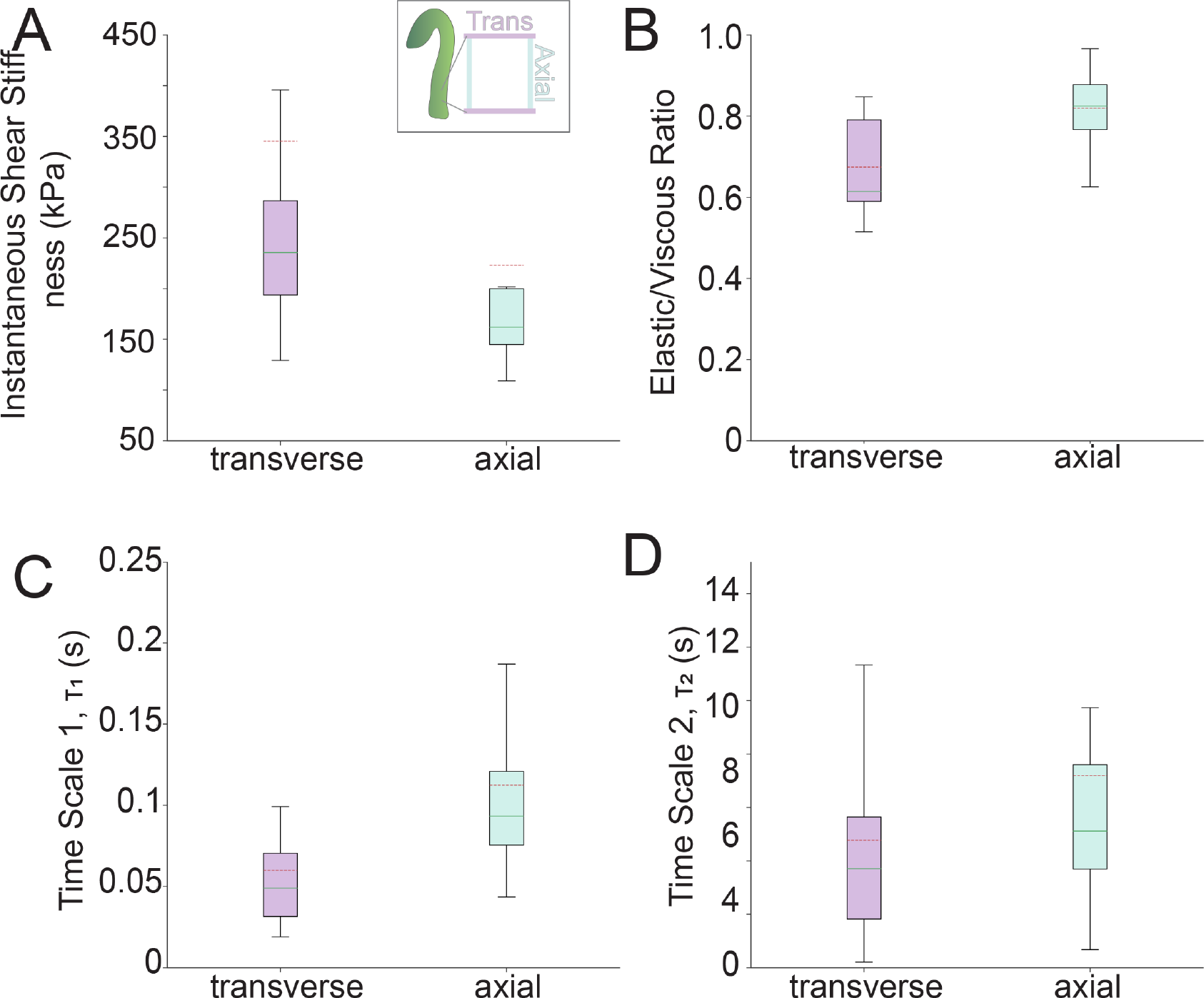
Plant model fit results – continuous line represents median of data and dashed line represents mean of data. a) ISS, or equivalent shear stiffness at time t=0, as found by the SLS2 model for both *Arabidopsis* wall types tested. b) Elastic/Viscous ratio as found by the SLS2 model for both *Arabidopsis* wall types tested. c) Time scale 1 of the SLS2 model for fitted to both *Arabidopsis* wall types tested. d) Time scale 2 of the SLS2 model for fitted to both *Arabidopsis* wall types tested.

In the pectin gels examined earlier, the time for relaxation did not change with DM. In the primary cell wall, we found that increasing DM increased both time-scales *τ*_1_ and *τ*_2_. Figures 4c and 4d show the time scales *τ*_1_ and *τ*_2_ as fitted by the SLS2 model. The mean values of both time-scales were higher in the axial walls (0.11 ± 0.02 s vs. 0.06 ± 0.01 s for *τ*_1_, and 7.2 ± 1.7 s vs 4.7 ± 0.9 s for *τ*_2_, axis and transverse walls respectively). However, only the difference in *τ*_1_ was found to be statistically significant (*p* < 0.005 for *τ*_1_ compared to *p* ≅ 0.16 for *τ*_2_). This difference was mirrored in the elastic/viscous ratio of the two wall types (Figure 4b); unlike the pure pectin gels tested earlier (Figure 2c), the plant cell walls exhibited different elastic/viscous ratios (0.82 ± 0.02 for axial walls and 0.67 ±s 0.03 for transverse walls, *p* < 0.005). Since we knew that elasticity (ISS) changed with wall type, we could predict that viscosity changed alongside it as in gels except that it was exaggerated. Given the difference in *τ*, a component of viscosity, it seemed reasonable to conclude that the inconsistent E/V ratio was due in part to altered creep time.

Since our creep experiments on plant cell walls did not always reach plateaus during the maximum experimental time, we used our SLS2 parameters to simulate response of the creep moduli for axial and transverse walls to a step loading (Figure 5b; lines represent averaged parameters, shaded areas are +/−standard error). This simulation yielded a clear graphical explanation for the E/V ratio difference and provides insight into the difference in time-scales: the axial walls were less stiff instantaneously but plateaued at a similar time to the transverse walls. This would require a slower rate of creep and higher effective time-scale, reflected in the time-scale data. Both axial and transverse wall compliance then plateaued at similar values, reflected in the overlap of their standard-error compliance in Figure 5b. Equivalently, these similar plateau compliances can be inverted and thought of plateau shear stiffnesses (PSS), as plotted in Figure 5a. Compared to the 55% difference in ISS, there was only a 21% difference in the means of the two PSS values (PSS = 301 ± 53 kPa for transverse walls, PSS = 248 ± 31 kPa for axial walls); the PSS statistical significance was *p* = 0.18 (2 s.f.) reflecting greater overlap than the ISS. As this closeness in PSS was not observed across DMs in pure HG gels, it seems there must be a cell-wall-specific explanation. Given the small tip diameter (10 nm) the AFM results should be fairly localized. Therefore, one possibility is that other locally situated components of the cell wall play a role in setting the plateau shear stiffness of the axial and transverse walls, the mightily strong cellulose [59] being the most probable candidate. If longer range effects do come into play, it is possible that the axial and transverse walls are architecturally and geometrically coupled in such a way that is only mechanically realized at longer time-scales. For example, the cellulose fibres in the cell wall would be contiguous across cell faces which may provide such coupling.

**Figure 5:**
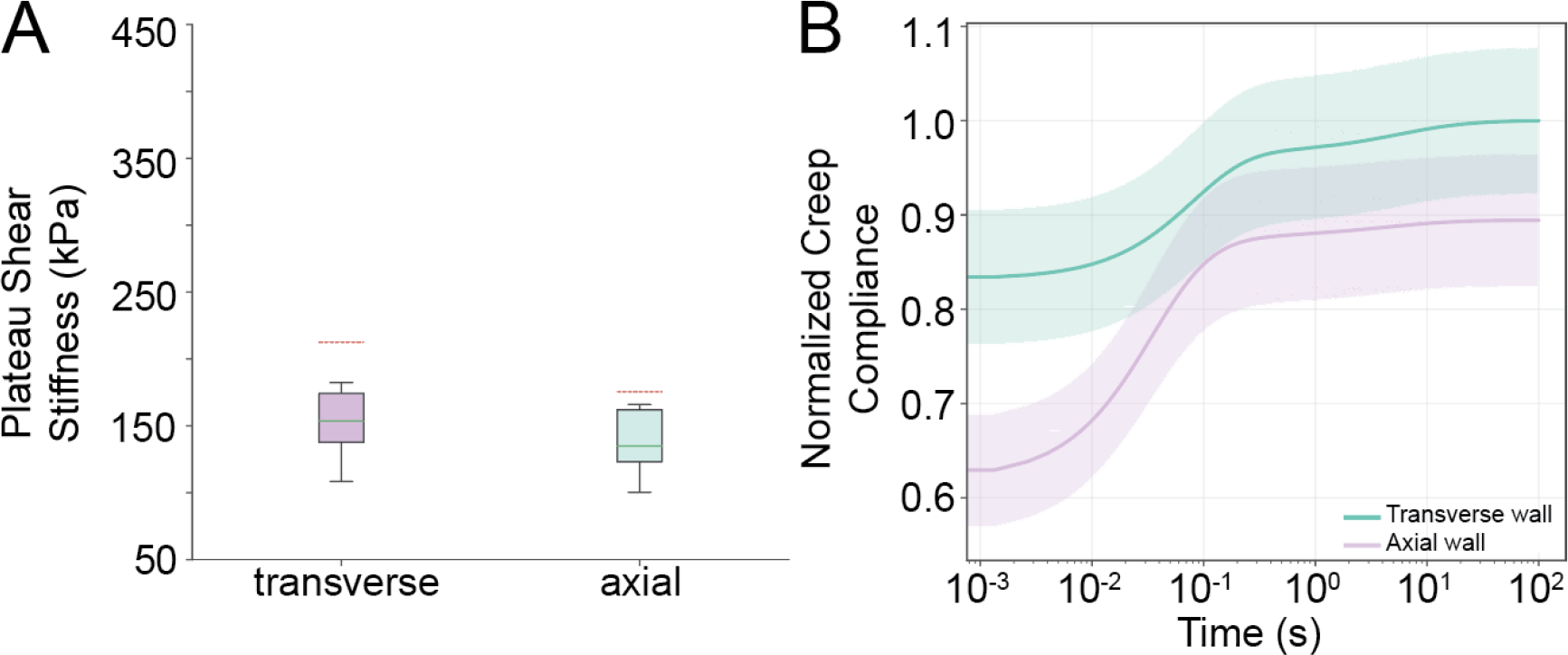
a) PSS, or equivalent shear stiffness as time *t* → ∞, as found by the SLS2 model for both *Arabidopsis* wall types tested – continuous line represents median of data and dashed line represents mean of data. b) Simulated creep compliance for both wall types predicted by the fitted SLS2 model subject to a step load.

## 4 Conclusions

In this study we developed an AFM-based viscoelastic methodology and used it to test the time-dependent mechanical properties of pectin gels and *Arabidopsis* hypocotyl cells. Linear viscoelasticity was identified as a suitable theoretical framework with which to analyse the time-dependent mechanical data from the HG gels. Two possible constitutive models were fitted and analysed for their suitability: an SLS2 and an FZ model. Both qualitative and quantitative evidence favoured the FZ model which could capture the power-law relaxation behaviour observed in HG gels. Through use of the above two models, it was observed that elasticity and viscosity changed in tandem with changes in DM; they were both negatively correlated with DM. From this, it was hypothesised that dynamic ionic bond reformation was the dominant mode of physical deformation during the force relaxation tests. The fitted models were then used to investigate several mixed DM HG gels. The non-linear relationship between gel strength and DM was found to explain the discrepancies between mixed and pure DM gels’ strengths. It should be noted that in our tests, the mechanical integrity of high DM gels (DM > 60) was negligible – they were unable to form coherent gels. Furthermore, even without increased gelation due to sugars, high DM pectin may play other roles such as permeability modification [60] and lubrication of other cell wall elements [61,62].

Viscoelastic testing of plant cell walls *in vivo* built upon our previous work on elasticity [17] and demonstrated that the SLS2 model provides a good fit to experimental creep data, in contrast to the gels where a fractional model was found to be superior. The instantaneous shear stiffness was found to be in line with indentation moduli found in elastic indentation tests. Both data sets were congruent with HG gel mechanical data here and the hypothesis that pectin DM is significantly correlated with short time-scale mechanical compliance. The time for creep to plateau was found to be slower in axial walls than transverse walls, in contrast to the gels which displayed similar timescales over the range of DMs tested. Lastly, the mean plateau shear stiffness was far closer between wall types than the instantaneous shear stiffness indicating the possibility of a mechanical coupling between axial and transverse walls that is only realised at longer time-scales. As cell walls must have anisotropic mechanical properties to allow anisotropic growth, the differences in stiffness at short time-scale but similarities at longer time-scale suggest that short time-scale mechanical properties may be the arbiters of growth processes. This would partly explain why elastic moduli have been successfully used as developmental correlates and in plant growth modelling [7, 17]. Given the fact that cell walls are clearly viscoelastic, the dynamic data presented here may be useful in development of more advanced growth models, computational or otherwise. Further work should be done to validate the SLS2 model as a constitute phenomenological model for plant cell walls so that fitting and prediction can be done, and to establish it as a standard that can be used to compare chemically treated plants or genetic mutants. In addition, if the SLS2 is fully validated, the physical significance of the 2 time scales should be established. Additional work could also explore longer time-scales, a finer spatial resolution, and probe the mechanisms that might be contributing to the apparent weak mechanical coupling between axial and transverse hypocotyl walls at longer time-scales.

## 5 Author Contributions

JLK participated in research project conception and direction, devised experimental methods, carried out AFM experiments of gels (apart from the two noted below), wrote code, performed all analyses, wrote manuscript, and prepared figures; TAT made gels and helped determine gel testing methods, performed initial elastic tests on the DM50(41/60) and DM50(33/70) gels, participated in discussions during project execution, and commented on manuscript; FB performed plant viscoelastic testing and commented on manuscript. SAB conceived of the project, consulted on experimental design, coordinated the project, secured funding, wrote the manuscript and arranged figures. All authors gave final approval for publication.

## 6 Acknowledgements

We would like to thank all members of the Braybrook Lab for helpful discussions during project execution. We would also like to thank Dr. Alessandra Bonfanti and Dr. Alexandre Kabla of the University of Cambridge Engineering Department for valuable discussions on fractional viscoelastic modelling, supporting viscoelastic theory and code implementation. We thank Herbsteith and Fox (DE) for their generous donation of pectin samples. The George and Lillian Schiff Foundation and Department of Engineering, University of Cambridge (UK) are thanked for their support of JLK.

---

**Table 1:** Mean Fractional Zener model derived parameters for all HG types +/− standard error.‘33/70’ is the DM41 mixture. ‘41/60’ and ‘33/70’ are the two DM50 mixtures.

| HG DM | ISS (kPa) | Springpot Parameter $\nu$ |
| --- | --- | --- |
| 33 | $128.9 \pm 23.7$ | $0.40 \pm 0.02$ |
| 41 | $73.4 \pm 5.9$ | $0.31 \pm 0.02$ |
| 50 | $11.4 \pm 0.8$ | $0.37 \pm 0.02$ |
| 33/50 | $67.7 \pm 1.2$ | $0.33 \pm 0.02$ |
| 41/60 | $40.5 \pm 2.8$ | $0.36 \pm 0.01$ |
| 33/70 | $52.4 \pm 6.5$ | $0.36 \pm 0.02$ |

**Table 2:**
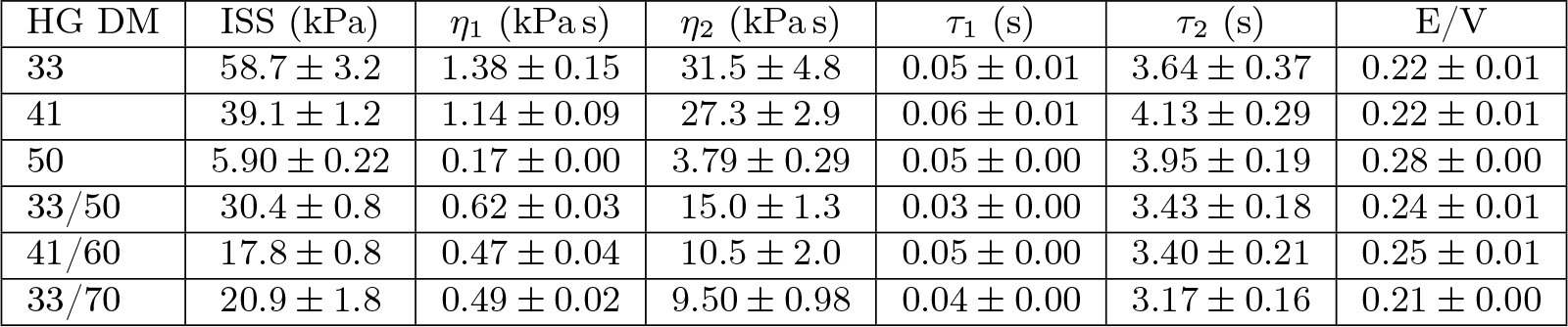
Mean SLS2 model derived parameters for all HG types +/−standard error. ‘E/V’ is Elastic/Viscous Ratio. ‘33/70’ is the DM41 mixture. ‘41/60’ and ‘33/70’ are the two DM50 mixtures.

| HG DM | ISS (kPa) | $\eta_1$ (kPa.s) | $\eta_2$ (kPa.s) | $\tau_1$ (s) | $\tau_2$ (s) | E/V |
| --- | --- | --- | --- | --- | --- | --- |
| 33 | $58.7 \pm 3.2$ | $1.38 \pm 0.15$ | $31.5 \pm 4.8$ | $0.05 \pm 0.01$ | $3.64 \pm 0.37$ | $0.22 \pm 0.01$ |
| 41 | $39.1 \pm 1.2$ | $1.14 \pm 0.09$ | $27.3 \pm 2.9$ | $0.06 \pm 0.01$ | $4.13 \pm 0.29$ | $0.22 \pm 0.01$ |
| 50 | $5.90 \pm 0.22$ | $0.17 \pm 0.00$ | $3.79 \pm 0.29$ | $0.05 \pm 0.00$ | $3.95 \pm 0.19$ | $0.28 \pm 0.00$ |
| 33/50 | $30.4 \pm 0.8$ | $0.62 \pm 0.03$ | $15.0 \pm 1.3$ | $0.03 \pm 0.00$ | $3.43 \pm 0.18$ | $0.24 \pm 0.01$ |
| 41/60 | $17.8 \pm 0.8$ | $0.47 \pm 0.04$ | $10.5 \pm 2.0$ | $0.05 \pm 0.00$ | $3.40 \pm 0.21$ | $0.25 \pm 0.01$ |
| 33/70 | $20.9 \pm 1.8$ | $0.49 \pm 0.02$ | $9.50 \pm 0.98$ | $0.04 \pm 0.00$ | $3.17 \pm 0.16$ | $0.21 \pm 0.00$ |

## Supplementary Information

### 1 Evaluation of Pectin Linear Viscoelasticity

#### 1.1 Theory

For a linear viscoelastic material, relaxation curves at different indentation depths will reduce to a single master curve when the following rescaling is applied

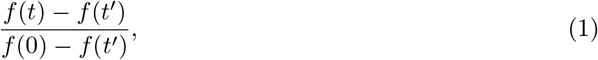

where in our case *t* = 0 was recalibrated to be the moment that the AFM reached its force setpoint and *t*′ was set to be 14 seconds after that, near the end of the relaxation hold. The poroelastic rescaling used was

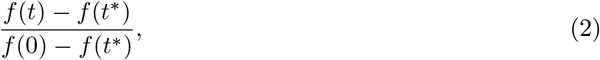

where *t\** = *αδ* as the tests did not generally reach a plateau within the time-frame of the experiment (see Moeendarbary et al. [1] and Strange et al. [2] for the more general case). The above curve is plotted against rescaled time 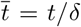. Also, as before, *t* = 0 was recalibrated to be the moment that the AFM reaches its force setpoint. *δ* was taken to be the average displacement occurring between 6 and 9 seconds into the experiment as a good approximation. The constant *α* was fixed for all curves and is arbitrary in the case of a poroelastic material that reduces to a master curve – though it will introduce a ‘pinch-point’ where all the curves meet. This can be seen algebraically by converting *t\** to rescaled time such that 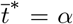 and noting that the numerator of Equation 2 will always be equal to 0 at this time point.

### 1.2 Results

During indentation of HG gels, there are likely several physical modes of deformation which occur simultaneously. These could include near-instantaneous extension and contraction of the pectin chain network, rearrangement of the network via breaking and reforming of ionic bonds, and flow of aqueous solution through the network. The dominant physical mechanisms observed depend on the pectin formulation, and the length and time-scales of the testing method used [2]. Various biochemical pectin formulations have been found to exhibit a range of deformation modes including linear and non-linear viscoelastic [3,4], and poroelastic [5]. Furthermore the algal-derived hydrogel alginate, which is comprised of a different polysaccharide to HG but also forms calcium cross-links [6], has been found to exhibit poroelastic or viscoelastic effects depending on whether the cross-linking mechanism used to form a gel was covalent or ionic respectively [7]. Thus, before a thorough comparison between different DM HG gels could be made, the appropriate theoretical framework to analyse the rheological data had to be identified.

First, a series of elastic indentation tests were conducted which enabled the determination of proxy forces which would yield desired indentation depths. This is required as AFM set-points must be specified in terms of force applied. DM41 HG gels were subjected to a series of rapid indentation tests across a range of forces from 500 to 6000 nN (approach speed = 50 µm s^−1^). The resulting force-indentation curves were analysed using the Hertzian contact model for a sphere in order to calculate the gels’ Young’s Moduli (E). This model was chosen over the DMT and JKR models due to the negligible adhesion observed in approach curves. These results also allowed assessment of linear elasticity. The change in mean average Young’s Modulus was only 18% over an order of magnitude force range which led us to conclude that, within the tested force range, the gels were behaving as linear elastic materials to a reasonable approximation (Figure 1a; E=58.9 ± 7.0 kPa, standard deviation; n=18, 3 points on 6 gels). The apparent change in E may also be increased by the lower signal/noise ratio at lower forces.

The time dependent behaviour of DM41 HG was then investigated in order to determine the relative importance of poroelasticity and viscoelasticity under AFM micro-indentation. AFM force relaxation tests, in which deformation was held constant while force was monitored over a prescribed time, were performed. Using the averaged force-displacement relationships found in the previous elastic experiment, 15 second relaxation tests were conducted on six DM41 gels at forces corresponding to 2 µm, 3 µm and 4 µm indentation depths (F = 1911 nN, 3185 nN and 4690 nN respectively). These forces corresponded well to the indentation depths predicted by our first set of experiments. To avoid any effects caused by pre-stressing, every relaxation test on a specific gel sample was done at a unique point. Suitable rescalings were then applied to the data to assess the suitability of a poroelastic or viscoelastic theoretical framework (See Methods, Equations 1 and 2). The poroelastic scaling did not collapse to a master curve except at the start and end (Figure 1c); the pinch-point at the end is due to the selection of the rescaling parameter, *α*. If the gels were poroelastic, the curves would collapse to a master-curve regardless of the value of *α*. However, this is clearly not the case, as corroborated by Figure 1d. Qualitatively, the linear viscoelastic rescaling resulted in a clear master-curve collapse (Figure 1b) suggesting it is the correct framework for subsequent analysis. A possible explanation for this can be constructed using scaling arguments. The experimental length scale here is 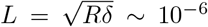. As *L*^3^ ≪ *L*^2^, the volume of water displacement may be less significant than the direct pressure applied onto the material by the indenter’s surface. Further, it may be that the permeability of the gel is high enough that there is negligible obstruction to fluid flow. A third explanation, similar to that proposed by Zhao et al. [7] for alginate relaxation with sufficient sample size, could be that the non-poroelastic behaviour is due to ionic cross-links reforming at a much faster rate than water migration. In summary, these data indicate that HG gels behave as linear-elastic viscoelastic solids under micro-indentation.

---

**Figure 1:**
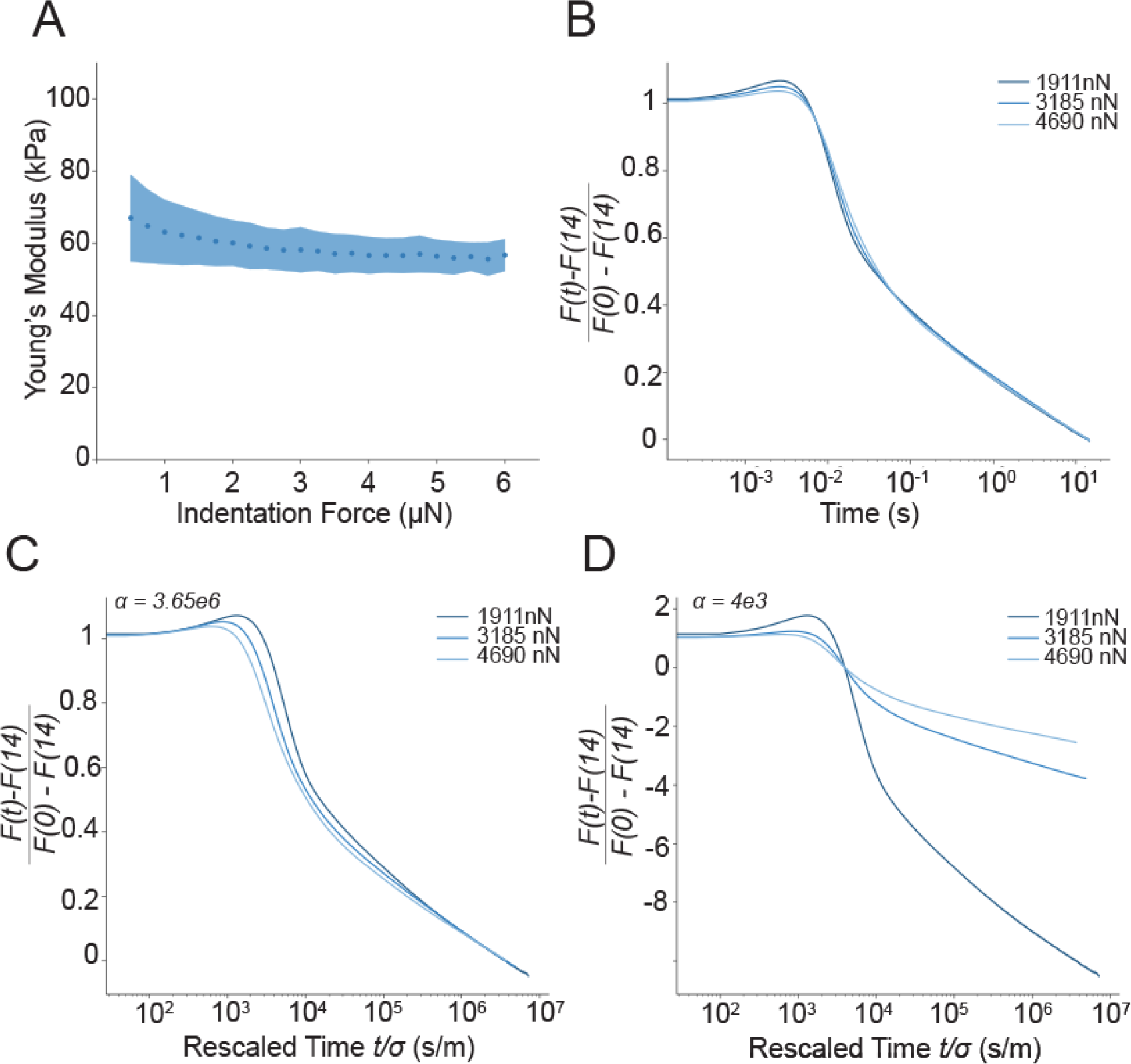
a) Young’s modulus vs. indentation force from elastics tests, DM41 HG gel. b) Linear viscoelastic rescaling of the averaged data from all DM41 viscoelastic tests at 3 forces corresponding approximately to 2 µm, 3 µm and 4 µm indentation depth. c) Poroelastic rescaling of the same DM41 averaged data as Figure 1b but with poroelastic rescaling and rescaling parameter *α* = 3.65*e*6. d) Poroelastic rescaling of the same DM41 averaged data as Figure 1b but with poroelastic rescaling and rescaling parameter *α* = 4*e*3.

